# Allosteric inhibition rescues hydrocephalus caused by catalytically inactive Shp2

**DOI:** 10.1101/2025.01.28.635289

**Authors:** Neoklis Makrides, Emily Sun, Hilal Mir, Ziyuan Jiang, Yihua Wu, Carlos Serra, Wellington V Cardoso, Neel H. Shah, Xin Zhang

**Author notes:** These authors contributed equally to this work (Neoklis Makrides, Emily Sun).

## Abstract

SHP2, a protein tyrosine phosphatase (PTP) crucial in Ras-MAPK signaling, is associated with various human congenital diseases and cancers. Here, we show that the catalytically inactive *Shp2^C459S^* mutation results in communicating hydrocephalus, similar to the catalytically activating *Shp2^E76K^* and *Mek1^DD^* mutants. Unlike previous mutants, however, *Shp2^C459S/+^* mutation uniquely affects ciliary development rather than neurogenesis, leading to reduced cilia density and impaired ciliary motility. Differential scanning fluorimetry revealed that *SHP2^C459S^*, SHP*2^E76K^* and *SHP2^C459S/E76K^* mutations all induce an open SHP2 conformation, but only *SHP2^C459S^* leads to aberrant GAB1 phosphorylation in cells expressing wild-type SHP2. This distinctive signaling pattern correlates with our observations in brain ventricular tissues of *Shp2^C459S/+^* mice, where Erk and Stat3 activities remain normal but Gab1 phosphorylation is elevated. Critically, we show that the hydrocephalus phenotype in *Shp2^C459S^* mice can be mitigated by allosteric inhibition of Shp2. These findings suggest that *Shp2*-associated hydrocephalus is driven by conformational changes rather than altered catalytic activity. Our results underscore the therapeutic potential of conformation-specific allosteric inhibitors in targeting both catalytically active and inactive *SHP2* mutants.

## Introduction

SHP2 (SH2-containing protein tyrosine phosphatase-2), a ubiquitously expressed enzyme encoded by the proto-oncogene *SHP2*, plays a critical role in various cell signaling pathways, including RAS/MAPK, PI3K/AKT and JAK/STAT signaling (Feng, 1999; Neel et al., 2003). The protein structure of SHP2 includes two N-terminal SH2 domains (N-SH2 and C-SH2), a protein tyrosine phosphatase (PTP) domain, and a C-terminal tail. At basal conditions, SHP2 exists in an autoinhibited state due to occlusion of the catalytic PTP cleft by the N-SH2 domain. This inhibition is overcome when phosphoproteins bind to the SH2 domains, prompting SHP2 to shift from a closed to an open, active configuration. Mutations in SHP2 are associated with developmental disorders such as Noonan syndrome (NS) and Noonan syndrome with multiple lentigines (NSML), as well as various cancers, including childhood leukemia (Gelb and Tartaglia, 2006). Gain-of-function mutations in NS disrupt the autoinhibitory mechanism, leading to constitutive activation of SHP2. Conversely, dominant-negative mutations in NSML impair the phosphatase activity of SHP2, resulting in reduced ERK signaling (Keilhack et al., 2005; O’Reilly et al., 2000). Despite these differences, the shared clinical features of NS and NSML, such as facial dysmorphism and cardiac abnormalities, suggest that SHP2 may operate through both catalytically dependent and independent mechanisms, though these pathways are not yet fully understood.

Hydrocephalus is a neurological condition characterized by an abnormal accumulation of cerebrospinal fluid (CSF) within the brain (Li et al., 2021). This condition can arise through various mechanisms such as blockages in the ventricular system, impaired fluid flow and absorption, or, less commonly, excessive CSF production. Previous studies have demonstrated that the neural expression of a gain-of-function mutation in the *Shp2* gene (*Shp2^E76K^*) can lead to hydrocephalus in mouse models (Zheng et al., 2018). The underlying pathological mechanism stems from the abnormal development of ventricular ependymal cell cilia, which was attributed to the reduced activation of Stat3 signaling. Notably, introducing a loss-of-function mutation (C459S) in the same *Shp2^E76K^* allele was shown to effectively prevent hydrocephalus, suggesting that aberrant Shp2 catalytic activity is responsible for reduced Stat3 signaling and subsequent ependymal cell abnormalities.

Cysteine 459, located within the protein tyrosine phosphatase (PTP) domain of Shp2, plays a crucial role in its enzymatic activity. In this study, we induced the expression of a single conditional *Shp2^C459S^*knock-in allele in mouse brain, which unexpectedly led to hydrocephalus resulting from abnormal ciliogenesis in the lateral ventricle ependymal cells. Mechanistically, we found that both the gain-of-function mutation E76K and the loss-of-function mutation C459S cause SHP2 to adopt an open conformation, but only C459S leads to elevated activation of GAB1 in the presence of the wild type SHP2. Furthermore, we showed that *Shp2^C459S^*-induced hydrocephalus could be ameliorated in vivo through allosteric inhibition of Shp2. These results indicate that the catalytic activity of *Shp2* is not essential for its role in hydrocephalus, highlighting the targeting of SHP2’s open conformation as a promising therapeutic strategy for catalytically inactive *SHP2*-related disorders.

## Results

### The Shp2 C459S mutation leads to communicating hydrocephalus

To investigate the role of Shp2 catalytic activity in hydrocephalus, we generate a *Shp2^C459S^* conditional allele containing a *loxP*-flanked stop cassette preceding the C459S point mutation in the *Shp2* locus (Fig. 1A) (Wang et al., 2024). The removal of the stop cassette in the germline of *Shp2^C459S/+^*mice with *EIIa-Cre* leads to embryonic lethality in the C57/BL6 background (n>20), in line with the autosomal dominant nature of *SHP2* mutations in humans. Subsequently, we induced the *Shp2^C459S^* in the brain using *Nestin-Cre*, which is specifically active in the neural and glial progenitors. This was validated using the *Ai9* Cre reporter line, which resulted in ubiquitous tdTomato reporter expression in the brain at birth (Fig. 1B), with the notable exception of the choroid plexus (Fig. 1C). Both *Nestin-Cre;Shp2^flox/flox^* and *Nestin-Cre;Shp2^flox/C459S^*mice exhibited early perinatal lethality, underscoring the crucial role of *Shp2* in brain development. In contrast, *Nestin-Cre;Shp2^C459S/+^* were born at the expected Mendelian ratio but developed signs of hydrocephalus by postnatal day 5 (P5), evident from their enlarged, domed skulls (Fig. 1D, arrowhead). Given the established role of Shp2 in Ras-MAPK signaling, we also induced a constitutively activated form of mitogen-activated protein kinase 1 (MEK1^DD^) in the brain using *Nestrin-Cre*. Notably, *Nestin-Cre;R26^MEK1DD^*pups also developed hydrocephalus, characterized by slight inflation of the brain and a transparent appearance in the visual cortex of both hemispheres indicating early cortical compression (Fig. 1E, arrowheads).

**Figure 1.**
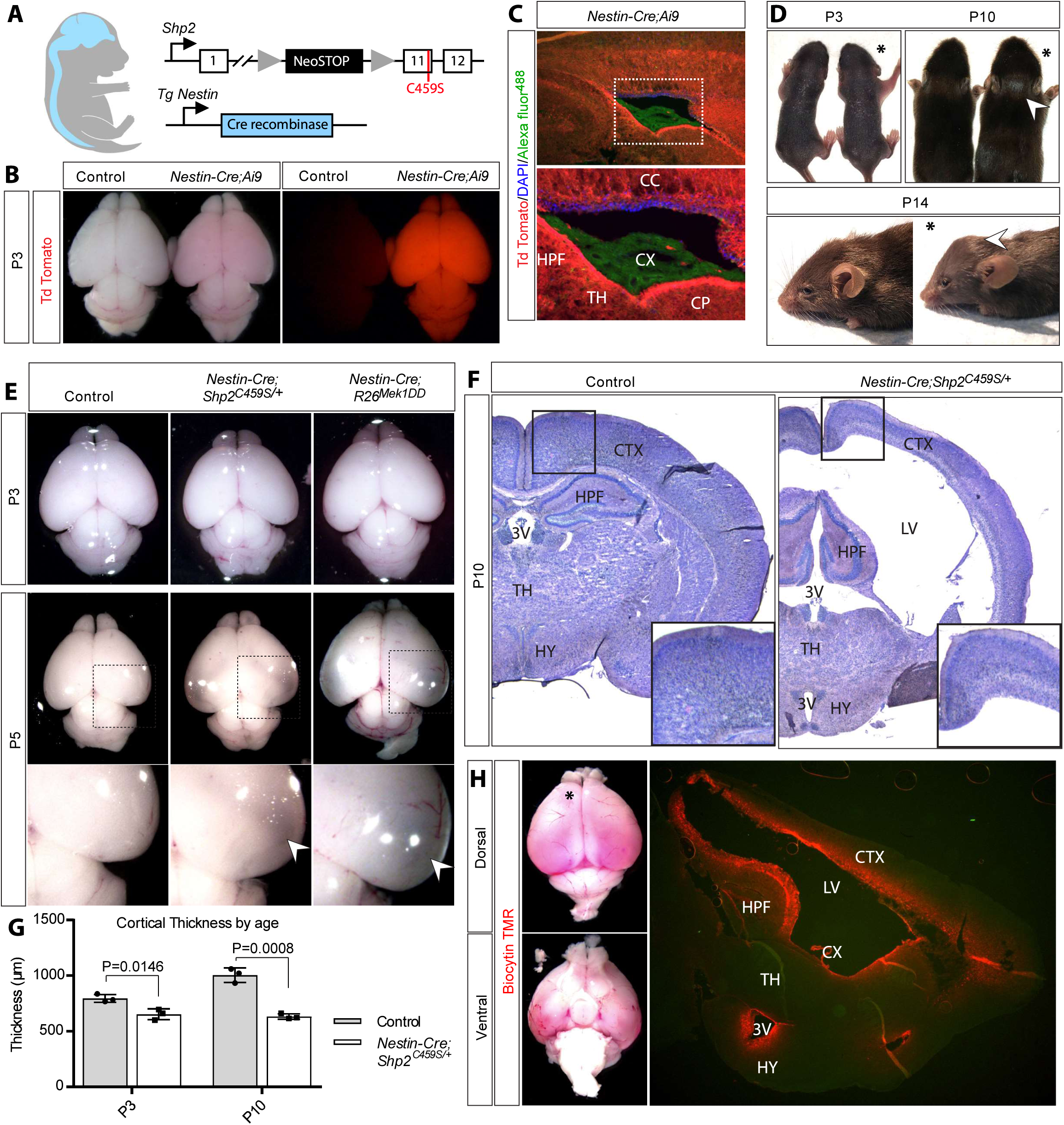
*Nestin-Cre;Shp2^C459S/+^* mice exhibit communicating hydrocephalus. **(A)** Schematic of the *Shp2^C459S^*conditional allele. **(B)** The efficiency of *Nestin-Cre* was assessed in neonatal brains using the Ai9 reporter under brightfield and epifluorescence microscopy. **(C)** The tdTomato reporter expression validated in sagittal cryosections of neonatal brains. **(D)** Phenotypic comparison of *Nestin-Cre;Shp2^C459S/+^* mice (asterisk) and control littermates rom P3 to P14.. **(E)** Brain morphology of wild type, *Nestin-Cre;Shp2^C459S/+^* and *Nestin-Cre;R26^MEK1DD^*mice. **(F-G)** Cortical thickness was measured in toluidine blue-stained coronal sections at bregma −1.22 mm, with data represented as mean ± S.D. of 4–5 biological replicates. **(H)** Alexa Fluor™ 594 Biocytin dye injected at the anterior horn of the lateral ventricle (asterisk) in P10 *Nestin-Cre;Shp2^C459S^* mice. Dye diffusion was analyzed in sagittal sections. Abbreviations: 3V- Third Ventricle, CC- Corpus Callosum, CP- Caudoputamen, CTX- Cortex, CX- Choroid Plexus, HPF- Hippocampus, HY- Hypothalamus, LV- Lateral Ventricle, TH- Thalamus.

To elucidate the nature of the hydrocephalus in *Nestin-Cre;Shp2^C459S/+^* mice, we conducted a histological assessment of the ventricular system on consecutive brain coronal sections stained with toluidine blue. This revealed significant ventricular inflation and progressive cortical thinning (Fig. 1F-G). Importantly, no blockages were detected in the ventricular pathway of the mutant mice, suggesting that the observed hydrocephalus was likely communicating hydrocephalus. This was confirmed by injecting biocytin dye into the anterior horn of the left lateral ventricle, which led to dye distribution throughout the entire ventricular system shown by sagittal sections (Fig. 1H), indicating unobstructed but poorly circulated cerebrospinal fluid (CSF) flow. These findings demonstrate that the expression of *Shp2^C459S^* in the brain causes communicating hydrocephalus.

### The Shp2 C459S hydrocephalic mice exhibit intact neural differentiation with impaired ciliogenesis

Genetic ablation of *Shp2* has been previously shown to disrupt the neurogenic to gliogenic switch during brain development (Zheng *et al*., 2018). However, the expression of neuronal marker NeuN and glial marker GFAP was similar between *Nestin-Cre;Shp2^C459S/+^* mutants and wild-type mice, which are in striking contrast to the significant reduction in NeuN and increased GFAP expression observed in *Nestin-Cre;R26^MEK1DD^*mice (Fig. 2A-B). Since decreased CSF flow due to malfunctioning ependymal cilia can lead to hydrocephalus, we further evaluated the integrity of the ventricular ependymal cells and their cilia in *Nestin-Cre;Shp2^C459S/+^* mice. Sox2 and Vimentin staining revealed that the ependymal cell lining remained intact throughout the ventricular system, particularly in the lateral and third ventricles (3V), subcommissural organ (SCO), and cerebral aqueduct (Aq)—regions frequently associated with hydrocephalus (Fig. 2C). Furthermore, we examined ependymal cell morphology using whole-mount F-actin staining of lateral ventricular walls (Fig. 2D). Unlike the disorganized apical lattice pattern seen in *Nestin-Cre;R26^MEK1DD^* mice, the ependymal cells in *Nestin-Cre;Shp2^C459S/+^* mice displayed a regular polygonal morphology, similar to that of wild-type controls (Fig. 2E). This indicates that the *Shp2^C459S^* mutation does not resemble either the *Shp2* null or gain-of-function *Mek1DD* mutant in disrupting the differentiation of ependymal cells from radial glial progenitors.

**Figure 2.**
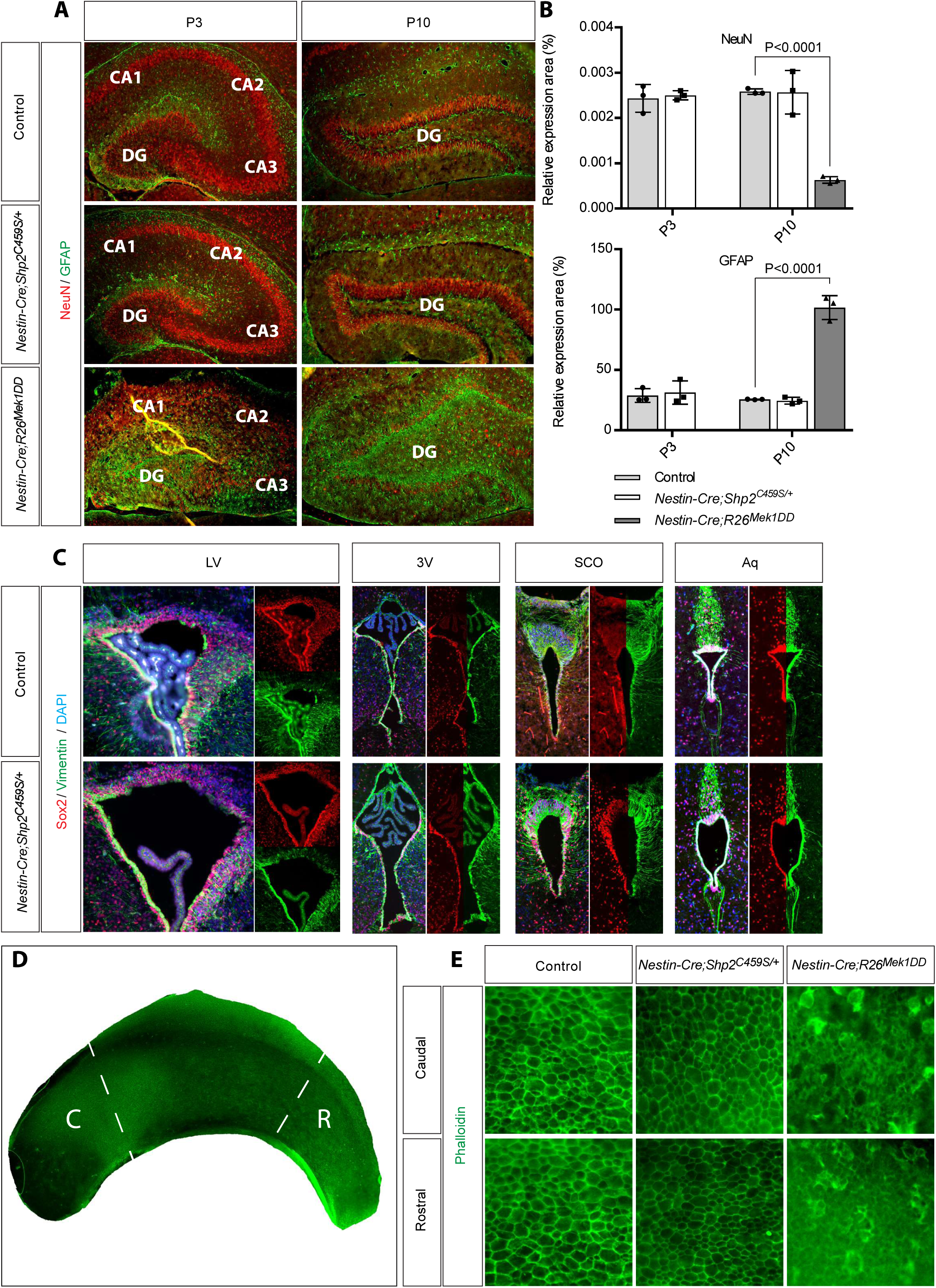
*Nestin-Cre;Shp2^C459S/+^* mice do not exhibit neural differentiation defects. **(A-B)** Hippocampal coronal sections of P3 and P10 brains harvested from wild type, *Nestin-Cre;Shp2^C459S/+^* and *Nestin-Cre;R26^MEK1DD^* mice stained with NeuN and GFAP. Quantification of NeuN-positive nuclei and GFAP expression presented as mean ± S.D. of 3 biological replicates. **(C)** P3 coronal sections of wild-type and *Nestin-Cre;Shp2^C459S^* brains stained with Vimentin and Sox2. **(D-E)** Lateral wall ependymal cell whole mounts of P3 brains stained with Phalloidin. Abbreviations: 3V- Third Ventricle, Aq- Cerebral Aqueduct, DG- Dentate Gyrus, LV- Lateral Ventricle, SCO- Subcommissural Organ.

We next assessed the structure and function of cilia in neonatal pups prior to the onset of hydrocephalus to eliminate confounding effects from increased ventricular pressure. β-IV Tubulin and Sox2 staining of lateral ventricle sections revealed a moderate reduction in cilia density on the medial wall and a complete absence on the lateral wall in *Nestin-Cre;Shp2^C459S/+^*mice (Fig. 3 A-C), while the cilia length remained unchanged (Fig. 3D). This is consistent with the expression of the transcription factor FoxJ1, a critical regulator of motile cilia formation, which was nearly absent in lateral wall sections of both *Nestin-Cre;Shp2^C459S/+^*and *Nestin-Cre;R26^MEK1DD^* mutant mice when compared to the wild type mice at P0 (Fig. 3E-F, arrowheads). Lastly, we examined cilia mobility in explant culture using high-speed live imaging, revealing that the ciliary beat frequency (CBF) was significantly lower in *Nestin-Cre;Shp2^C459S/+^*mutants (Fig. 3G-H). These results suggest that *Shp2^C459S^* mutation reduced the number of ciliated cells and their mobility, which leads to disruption of cerebrospinal fluid (CSF) flow and hydrocephalus.

**Figure 3.**
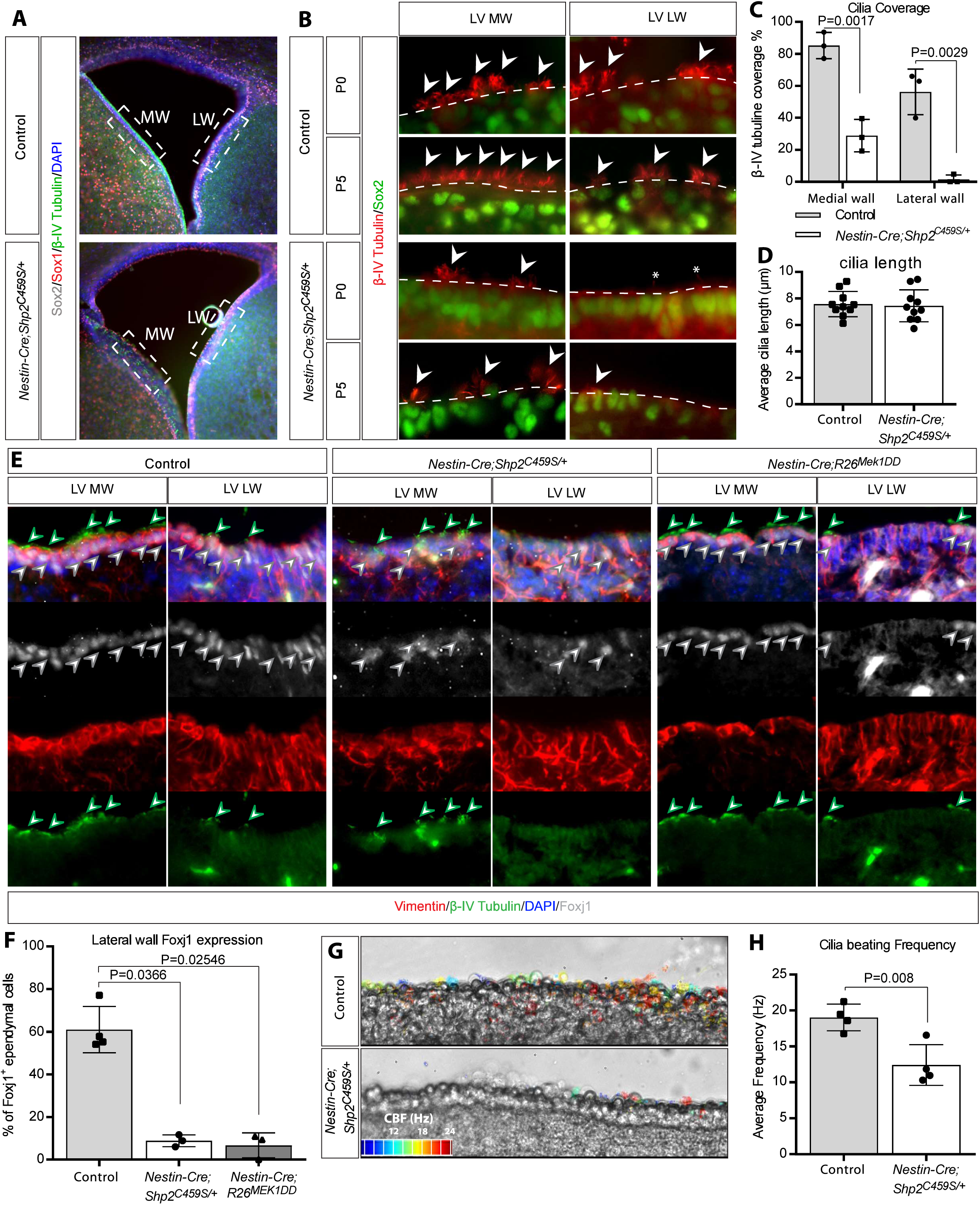
SHP2^C459S^ impairs ciliogenesis. **(A)** P0 coronal sections from wild-type and *Nestin-Cre;Shp2^C459S/+^* brains stained with Sox1, Sox2 and β-IV Tubulin. **(B)** Cilia density examined on lateral (LW) and medial (MW) ventricular walls. **(C)** Quantification of ciliary coverage based on β-IV Tubulin staining. **(D)** Average cilia length from medial wall sections represented as mean ± S.D. of 10 biological replicates. **(E-F)** Coronal lateral ventricle sections of neonatal wild-type, *Nestin-Cre;Shp2^C459S/+^* and *Nestin-Cre;R26^MEK1DD^* brains stained with Foxj1, Vimentin, and β-IV Tubulin. **(G-H)** High-speed video microscopy of lateral wall cilia from P3 coronal sections. Ciliary beating frequency (CBF) shown as heatmaps and plots (mean ± S.D. of 3 biological replicates).

Given that ciliogenesis in lateral wall ependymal cells progresses from the caudal to the rostral end during the perinatal period, we analyzed cilia density in lateral wall whole mounts to avoid potential sectioning artifacts. This analysis showed significantly fewer ciliated cells in the caudal portion of the *Nestin-Cre;Shp2^C459S/+^*lateral walls compared to controls (Fig. 4A-B). Moreover, the ciliogenesis front in these mutants was delayed by approximately 20% (Fig. 4C-D), indicating a maturation defect. This finding is important as ciliogenesis was normal in *Nestin-Cre;Shp2^flox/flox^* pups, as evidenced by the presence of ciliated cells in the rostral lateral walls (Fig. 4A-B). Conversely, although *Nestin-Cre;R26^MEK1DD^* mutants also exhibited reduced cilia density, their ciliogenesis front was not affected. These observations suggest that the phenotype seen in *Nestin-Cre;Shp2^C459S/+^* mutants may be independent of ERK/MAPK signaling. This conclusion is further supported by the combined phenotype of delayed ciliogenesis and ependymal cell disorganization in the *Nestin-Cre;Shp2^C459S/+^;R26^MEK1DD^*double mutants, indicating that activation of ERK signaling fails to rescue *Shp2^C459S^* mutation.

**Figure 4.**
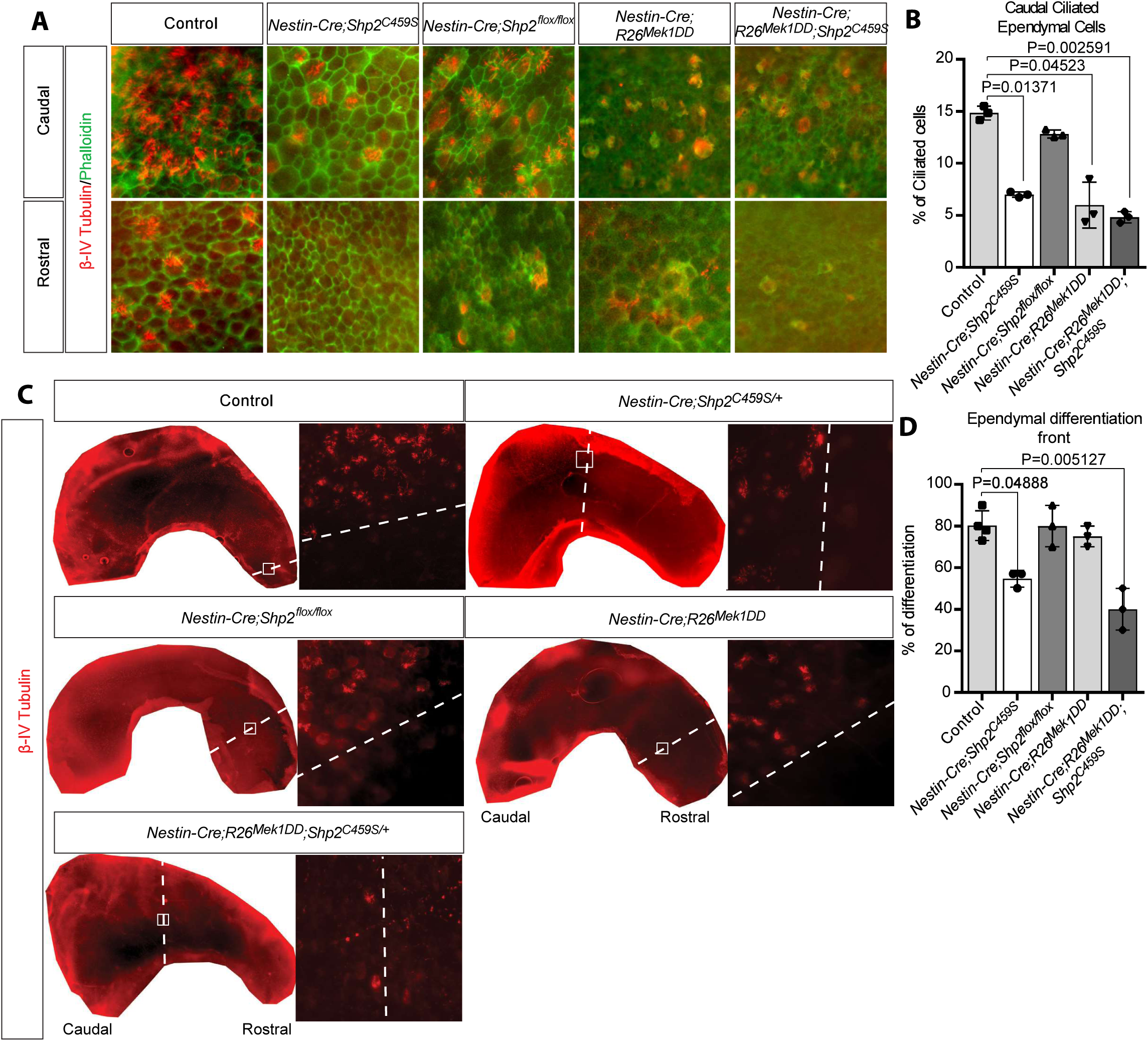
SHP2^C459S^ causes a delay in lateral wall ciliogenesis. **(A-B)** Lateral wall ependymal cell whole mounts of neonatal wild-type, *Nestin-Cre;Shp2^C459S/+^*, *Nestin-Cre;Shp2^flox/flox^*, *Nestin-Cre;R26^MEK1DD^* brains stained with Phalloidin and β-IV Tubulin. **(C-D)** Ciliogenic front assessed in lateral wall flat mounts. Proportions of ciliated ependymal cells at the caudal region and ciliogenic fronts presented as mean ± S.D. of 3 biological replicates.

### *SHP2* C459S mutation causes aberrant GAB1 activation

Previous studies have shown that the hyperactivating E76K mutation in *SHP2* leads to hydrocephalus, which can be rescued by the introduction of an additional C459S mutation (Zheng *et al*., 2018). Since both the E76K and C459S mutations are known to promote the open conformation of *SHP2*, it raises the question of whether the double mutant (C459S/E76K) might shift toward the closed conformation (Padua et al., 2018; Sha et al., 2023). To explore this possibility, we performed differential scanning fluorimetry to assess the open-closed conformational equilibria of the *SHP2* mutants through thermal melt-temperature analysis (Serbina and Bishop, 2023). Interestingly, the thermal melt-temperature curves of both the single C459S mutant and the double C459S/E76K mutant closely resembled that of the previously characterized open-conformation E76K mutant (Fig. 5A). In contrast, the wild-type protein displayed greater thermal stability, requiring elevated temperatures for denaturation. These findings suggest that the rescue of hydrocephalus in the C459S/E76K double mutant is unlikely to arise from the adoption of a closed conformation.

**Figure 5.**
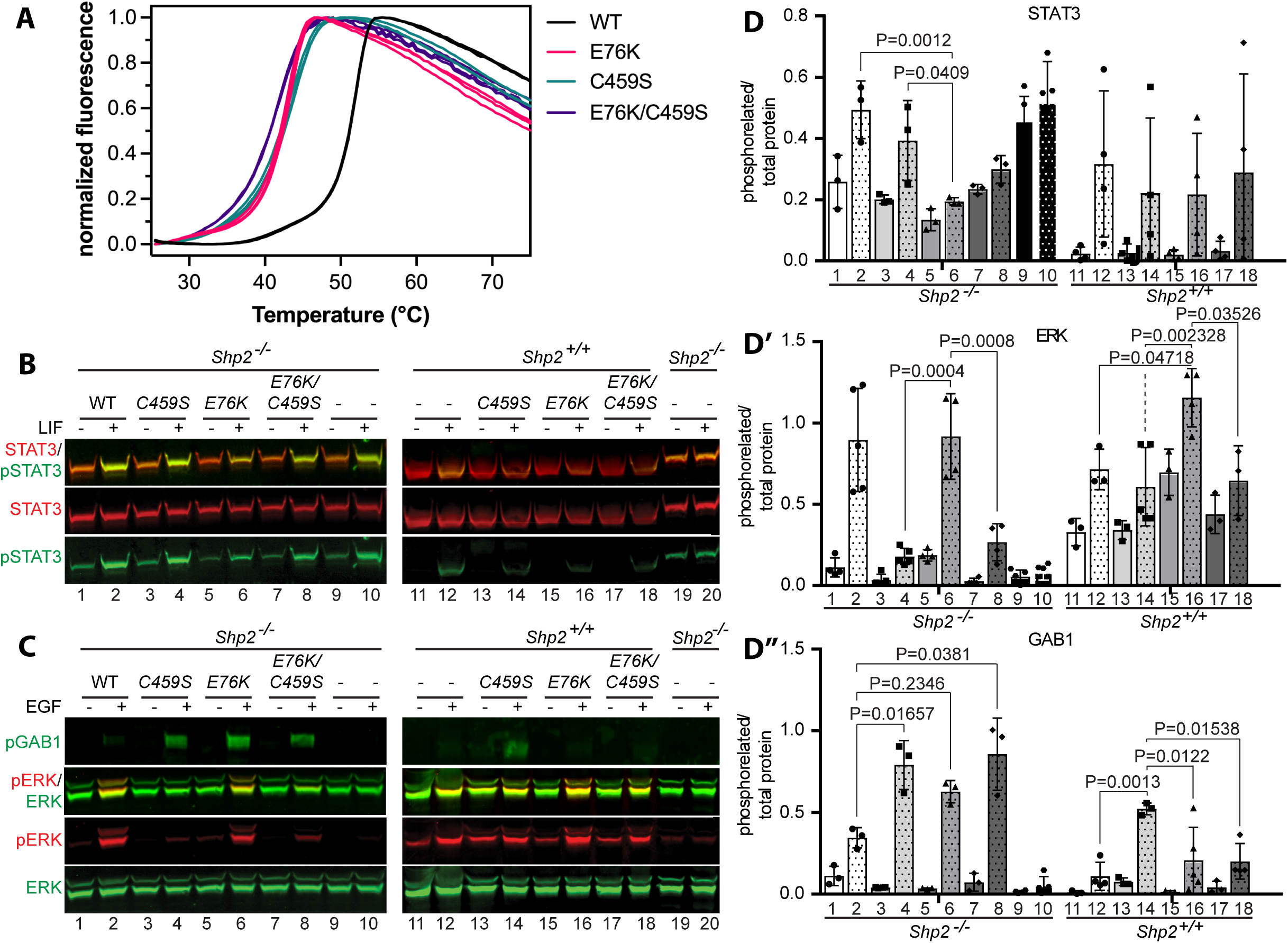
Open conformation of SHP2 leads to elevated pGAB1 levels. **(A)** Thermal denaturation curves of wild-type SHP2, SHP2^E76K^, SHP2^C459S^ and SHP2^E76K/C459S^. **(B-C)** Immunoblot of pGAB1, pSTAT3 and pERK of cell lysates from SHP2^−/−^ HEK293T and wild type HEK293T cells transfected with wild type SHP2, SHP2^E76K^, SHP2^C459S^ or SHP2^E76K/C459S^ and treated with either LIF (B) or EGF (C). **(D-D’’’)** Phosphorylated protein intensity relative to total protein shown as mean ± S.D. of 3 biological replicates.

To further investigate how these *SHP2* variations affect cell signaling, we performed in vitro experiments using wild-type and *SHP2*-null HEK293T cells transfected with various *SHP2* point mutants. The negative regulatory role of SHP2 in the JAK/STAT signaling cascade was assessed by treating the cells with LIF. As previously reported, *SHP2*-null cells displayed elevated basal pSTAT3 levels compared to wild-type HEK293T cells (Fig. 5B and D). Notably, cells expressing the E76K variant showed markedly decreased pSTAT3 levels, both at baseline and following LIF stimulation, when compared to cells expressing wild-type *SHP2*. In contrast, neither the C459S mutant nor the E76K/C459S double mutant altered pSTAT3 levels significantly. These results demonstrate that, unlike the gain-of-function E76K mutation, the C459S mutation does not affect STAT signaling.

We next examined the positive regulatory role of SHP2 in MAPK signaling by stimulating cells with EGF. As expected, *SHP2*-deficient cells exhibited minimal pERK activation in response to EGF, which was restored by transfection with wild-type *SHP2* (Fig. 5C and D’). The E76K *SHP2* mutant promoted ERK signaling comparable to wild-type *SHP2* following EGF stimulation, albeit with elevated basal pERK levels, consistent with this mutation’s constitutive activity. Interestingly, transfection with the C459S *SHP2* mutant also modestly restored ERK1/2 phosphorylation. In contrast to pERK, pAKT was unchanged across all SHP2 variants (C459S, E76K, and E76K/C459S), but pGAB1 was markedly elevated compared to wild-type SHP2 (Fig. 5C, D’’ and D’’’). In wild-type HEK293T cells, which better reflect patient genetics where mutations occur alongside a wild-type allele, the E76K mutant induced the highest levels of pERK following EGF stimulation, while C459S and E76K/C459S showed similar pERK activation as wild type. Interestingly, while pGAB1 levels remained markedly elevated in cells transfected with the C459S mutant, the increase was far less pronounced in cells transfected with the E76K or E76K/C459S mutants. These results demonstrate that, in the presence of wild type *SHP2*, the C459S *SHP2* mutation specifically activates GAB1 signaling without affecting ERK and AKT signaling.

### Allosteric inhibition of SHP2 ameliorates hydrocephalus in the C459S mutant mice

To validate our in vitro findings, we analyzed signaling dynamics in the lateral ventricular wall through western blotting and immunostaining. *Nestin-Cre;R26^MEK1DD^*mice exhibited enhanced pERK signaling from P0 onward, followed by increased pStat3 levels at P7 (Fig. 6A-C). By contrast, *Nestin-Cre;Shp2^C459S/+^*mice showed normal pErk and pStat3 levels at P0, with elevations emerging only at P7. In both mutants, there was no appreciable change in pAkt level. The delayed activation pattern in *Nestin-Cre;Shp2^C459S/+^*mice suggests that elevated pErk and pStat3 likely reflect a secondary neuroinflammatory response rather than direct pathway activation. Significantly, analysis of lateral wall lysates revealed that Gab1 phosphorylation was consistently elevated in *Nestin-Cre;Shp2^C459S/+^*mice as early as embryonic day 15 (E15), preceding any changes in ERK signaling (Fig. 5D-E). These temporal dynamics corroborate our in vitro observations and suggest Gab1 hyperphosphorylation as an initiating event in *Shp2^C459S^*hydrocephalus mice.

**Figure 6.**
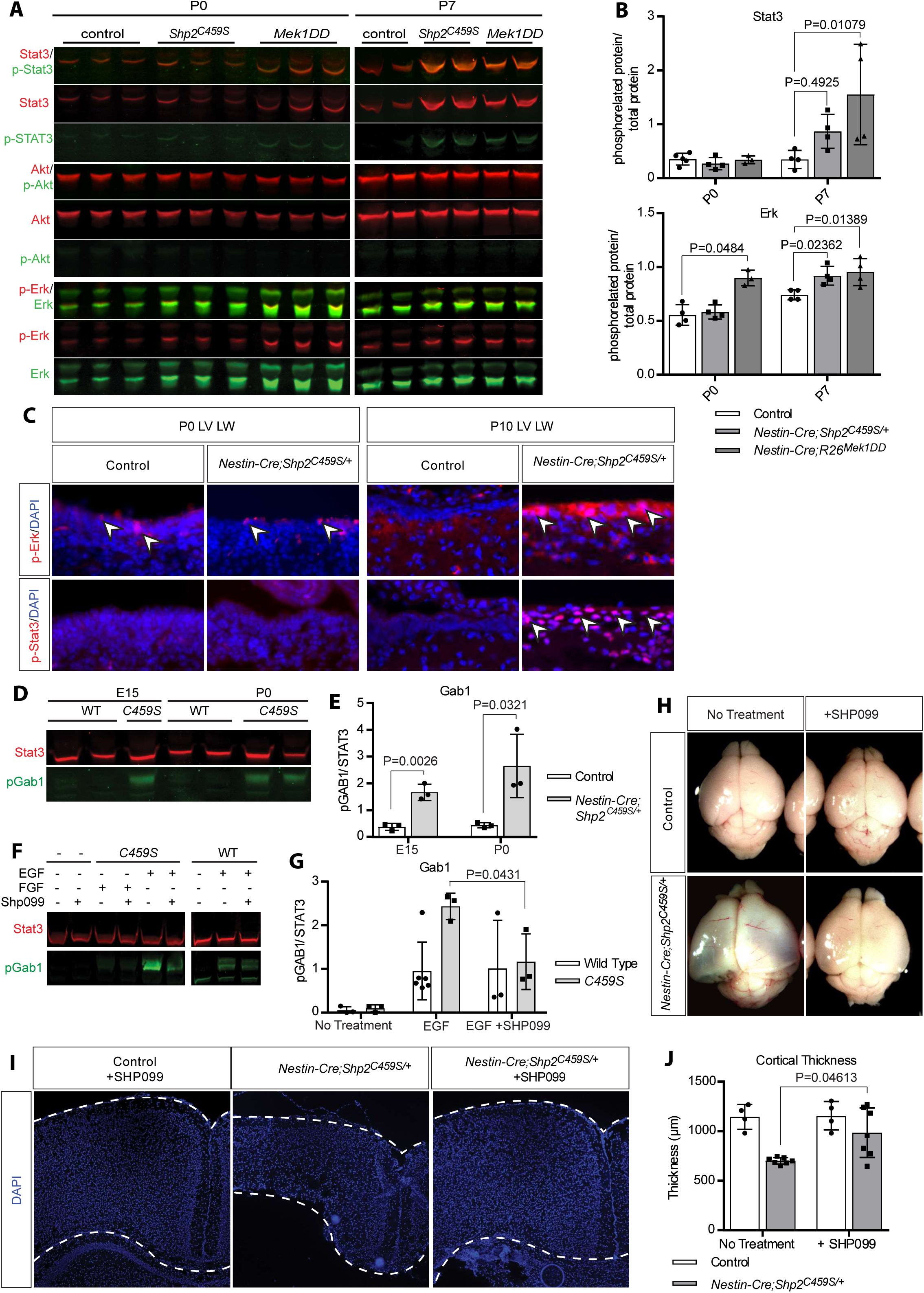
Allosteric inhibition of SHP2 reduces the severity of the *Nestin-Cre;Shp2^C459S/+^* hydrocephalus. **(A-B)** Immunoblots of P0 and P7 lateral wall lysates probed for pSTAT3, pAKT, and pERK. Phosphorylation levels relative to total protein represented as mean ± S.D. of 3 biological replicates. **(C)** pERK and pSTAT3 activation assessed in P0 and P7 lateral wall sections via immunostaining. **(D-E)** Immunoblots of E15 and P0 lateral wall lysates probed for pGAB1 with STAT3 as the loading control. **(F-G)** Immunoblot of pGAB1 and pSTAT3 of SHP2^−/−^ HEK293T cell lysates transfected with wild type SHP2 or SHP2^C459S^ and treated with either EGF or FGF ± SHP099 inhibitor. pGAB1 levels relative to total STAT3 shown as mean ± S.D. of 3 biological replicates. **(H)** Brain morphology of wild type and *Nestin-Cre;Shp2^C459S/+^* P10 pups with or without SHP099 injections at P0. **(I-J)** The cortical thickness of *Nestin-Cre;Shp2^C459S/+^* and control littermates ± SHP099 at P10, assessed in DAPI-stained coronal sections at bregma −1.22 mm. Data presented as mean ± S.D. of 6–17 biological replicates.

To explore potential therapeutic approaches, we tested whether the pharmacological inhibition of SHP2 could suppress C459S-induced GAB1 signaling. In wild-type HEK293T cells transfected with the C459S mutant, treatment with the allosteric SHP2 inhibitor SHP099 significantly reduced pGAB1 levels (Fig. 6F-G). Importantly, this effect was specific to the C459S mutation, as SHP099 did not alter pGAB1 levels in cells expressing only wild-type *SHP2*. Encouraged by these results, we evaluated SHP099’s therapeutic potential in *Nestin-Cre;Shp2^C459S/+^*mice. Intraperitoneal administration of SHP099 to newborn pups significantly reduced bilateral ventricular enlargement when assessed at P10, while having no adverse effects on control animals (Fig. 6H-L). Histological examination confirmed consistent cortical preservation in SHP099-treated mutant mice. These results establish allosteric SHP2 inhibition as a promising therapeutic approach for hydrocephalus arising from catalytically inactive *SHP2* mutations.

## Discussion

The central role of SHP2 in receptor tyrosine kinase (RTK) signal transduction is underscored by its involvement in diverse human pathologies, including multiple forms of leukemia, Noonan syndrome (NS), and Noonan syndrome with multiple lentigines (NSML) (Gelb and Tartaglia, 2006). Despite its biological significance, the mechanism by which this tyrosine phosphatase positively regulates RTK signaling remains unclear. Adding to this complexity is the phenotypic similarities between loss-of-function and gain-of-function *SHP2* mutations. Here, we report that the catalytically inactivating C459S mutation in *Shp2* induces communicating hydrocephalus in mice. Our findings show that the C459S mutation does not phenocopy either loss-of-function *Shp2* null mutations or gain-of-function *Mek* mutations in brain development or MAPK signaling. Instead, the mutation uniquely activates Gab1 signaling in the presence of wild-type Shp2. Importantly, treatment with an allosteric SHP2 inhibitor mitigated hydrocephalus in these mice. Although allosteric inhibitors are in general designed to target enzymatic activity, SHP2 allosteric inhibitors stabilize its closed state, which could also impact binding and scaffolding function. Thus, our results provide proof of principle that such inhibitors can also prevent the deleterious effects caused by some catalytically inactive mutations. These findings expand the therapeutic potential of SHP2 inhibitors and provide insights into the unique biological activity of the C459S mutation.

Our study revealed that the catalytically inactivating C459S mutation induces communicating hydrocephalus through impaired ciliogenic maturation of the lateral ventricles, rather than disrupted neural differentiation. This delay in ependymal cell ciliogenesis was unique to *Nestin-Cre;Shp2^C459S/+^* mutants and absent in *NesCre;Shp2^flox/flox^* mice, indicating that the defect likely does not arise from loss of Shp2 function. While premature activation of the gliogenic switch was observed in both *Nestin-Cre;R26^MEK1DD^* mice and mice expressing the catalytically activating SHP2 E76K mutation (Zheng *et al*., 2018), this phenotype was not present in *Nestin-Cre;Shp2^C459S/+^* mutants. This distinction suggests that the C459S mutation operates independently of ERK/MAPK signaling. Further supporting this conclusion, *Nestin-Cre;Shp2^C459S/+^* mice exhibited delayed ciliogenic front progression, whereas *Nestin-Cre;R26^MEK1DD^*mice showed an overall reduction in ciliated cells throughout the ventricle. Moreover, unlike the SHP2 E76K mutant, pStat3 levels during ependymal cell ciliogenesis remained comparable between wild-type and *Nestin-Cre;Shp2^C459S/+^* mutant mice. Collectively, these findings suggest that the hydrocephalus phenotype in *Nestin-Cre;Shp2^C459S/+^* mutant mice mutants develops independently of both MAPK and STAT pathway dysregulation.

Our biochemical analysis confirmed the in vivo distinction among the *SHP2* variants. In *SHP2*-null cells, the C459S mutant failed to restore EGF-stimulated pERK activation, while the E76K variant maintained elevated basal pERK levels. Despite its known substrate-trapping properties, the C459S mutant did not alter pERK or pSTAT3 levels when expressed in wild-type cells. Interestingly, differential scanning fluorimetry analysis revealed that all three mutations - C459S, E76K, and the C459S/E76K double mutant - promote an open conformation of the enzyme. Previous studies have shown that Gab1 and Shp2 act as partners in liver regeneration, while Gab2 is required for myeloid transformation induced by the *E76K Shp2* mutation (Bard-Chapeau et al., 2006; Mohi et al., 2005). Notably, the *C459S*, *E76K*, and *C459S/E76K* mutations caused aberrant GAB1 phosphorylation in *SHP2*-null cells. However, in wild-type cells, only the *C459S* variant significantly elevated pGAB1 levels. GAB proteins function as both docking partners for the SH2 domains of SHP2, as well as dephosphorylation substrates of the phosphatase domain (Montagner et al., 2005). Like the activating E76K mutation, the C459S mutation stabilizes an open conformation of SHP2 that is highly competent for binding GAB1, but unlike E76K, C459S is catalytically inactive and cannot dephosphorylate GAB1. Given this, we propose that the *C459S* mutant binds to phosphorylated GAB1, thereby protecting it from dephosphorylation by wild-type SHP2.

The allosteric SHP2 inhibitor SHP099 was originally developed to inhibit the catalytic activity of wild type SHP2 by stabilizing it in a closed, inactive conformation (Chen et al., 2016). Subsequent studies revealed that SHP099 could also force the constitutively active *E76K* variant into a closed state, though higher drug concentrations were required (LaRochelle et al., 2018; Padua *et al*., 2018). Since both *E76K* and *C459S* mutants favor an open conformation, we hypothesized that SHP099 might similarly suppress the pathological effects of the catalytically inactive *C459S* variant. Remarkably, we demonstrated that SHP099 effectively inhibited GAB1 hyperphosphorylation caused by the *C459S* mutant in cellular studies. More importantly, SHP099 treatment significantly ameliorated hydrocephalus in mice expressing the *C459S* mutant in the brain. These results reveal that SHP2’s pathological effects in hydrocephalus stem from its open conformation rather than its enzymatic activity—a finding that aligns with clinical observations of NSML syndrome arising from catalytically inactive SHP2 mutations. By demonstrating that allosteric inhibitors can counteract the pathological effects of catalytically inactive mutants, we establish a new paradigm for treating diseases caused by these previously “undruggable” SHP2 variants.

## Material and methods

### Mice husbandry

*Shp2^LSL^*^−*C459S*^ was previously generated as described (Wang *et al*., 2024). We obtained *Shp2^flox^*from Gen-sheng Feng (UCSD, San Diego, CA) (Zhang et al., 2004). Additional mouse lines used in this study were *EIIa-Cre* (Jackson Laboratory, RRID:IMSR_JAX: 003724), *Nestin-Cre* (Jackson Laboratory, RRID:IMSR_JAX: 003771), *R26^MEK1DD^*(Jackson Laboratory, RRID:MGI:012352), and *Ai9* (Jackson Laboratory, RRID:MGI:3817869). The mice were kept in a mixed genetic background and were housed at a constant temperature of 21°C with access to water and conventional chow diet ad libitum. All husbandry and experimental procedures complied to Columbia University’s Institutional Animal Care and Use Committee regulatory standards.

### Biocytin dye injection

Hydrocephalic P10 mice were deeply anesthetized with ketamine (150 mg/kg)/xylazine (15 mg/kg) and 5 μl of Alexa Fluor™ 594 Biocytin dye (A12922, Molecular Probes™) (1% in 0.9% saline) were slowly injected in the anterior horn of the lateral ventricle using a glass capillary needle (50 μm). The dye was allowed to diffuse throughout the ventricular system for 5 minutes prior to sacrificing the pups and extracting the brain.

### Immunohistochemistry and Toluidine blue staining

Mice were euthanized through decapitation and the brains were dissected out and fixed in 4% paraformaldehyde (PFA) overnight at 4°C. For paraffin sections, the tissues were dehydrated in EtOH and embedded in paraffin according to standard procedures. For cryosections, the brains were cryopreserved in 30% sucrose overnight at 4°C. The brains were embedded in cryomolds containing Tissue-Tek® optimal cutting temperature (OCT) compound (Sakura, Japan) and sectioned on a Leica CM1850 UV-3-1 Cryostat Microtome (Leica Biosystems, US) at 10 µm thickness. For immunostaining, antigen retrieval was performed on the sections using a heat steamer and 10 mM sodium citrate buffer (pH 6.0) prior to primary antibody staining (see Table 1) as previously described (Carbe et al., 2012; Carbe and Zhang, 2011). The slides were mounted with n-propyl gallate anti-fading reagent (P3130, Sigma-Aldrich) and imaged using a Leica DM5000-B fluorescence microscope. For toluidine blue staining, deparaffinized sections were covered with 0.3% toluidine blue dye for 1 minute. The sections were subsequently washed in Phosphate Buffer Solution (PBS) for 5 minutes three times and mounted with VectaMount AQ mounting medium (Vector Laboratories, LS-J1038-60).

**Table 1.** Antibodies used in the study.

| Species | Antibody | Identifier | Source |
| --- | --- | --- | --- |
| Mouse | Anti-AKT (clone 40D4) | Cat#2920 | Cell Signaling |
| Mouse | Anti-β IV Tubulin | Cat# ab11315 | Abcam |
| Rabbit | Anti- ERK1/2 (137F5) | Cat#4695 | Cell Signaling |
| Mouse | anti-FOXJ1 (clone 2A5) | Cat#14-9965-80 | Thermo Fisher Scientific |
| Rabbit | Anti-GFAP | Cat#0334 | Dako |
| Mouse | Anti-NeuN | Cat#mab377 | Chemicon |
|  | Phalloidin-488 | Cat# A12379 | Thermo Fisher Scientific |
| Rabbit | Anti-pAKT (473) | Cat#9271 | Cell Signaling |
| Rabbit | Anti-pERK1/2<br>(Thr202/Tyr204) | Cat#4370 | Cell Signaling |
| Rabbit | Anti-pGab1 | Cat#3233 | Cell Signaling |
| Rabbit | Anti-pStat3 (Tyr705) | Cat#9145 | Cell Signaling |
| Goat | Anti-Sox1 | Cat# AF3369-SP | R&D |
| Rat | Anti-Sox2 | Cat#14-9811-82 | Thermo Fisher Scientific |
| Mouse | Stat3 (clone 124H6) | Cat#9139 | Cell Signaling |
| Rabbit | Anti-Vimentin (D21H3) | Cat#5741 | Cell Signaling |

### Lateral wall whole mounts

Brains extracted from the mice were immediately placed in PBS and the lateral walls of the lateral ventricle were dissected as previously described (Mirzadeh et al., 2008). Briefly, the brains were split along the central sulcus and the occipital and olfactory cortex were dissected out. The hippocampus was subsequently removed using fine tip tweezers allowing for the separation of the ventricular medial and lateral walls. The dissected lateral walls were fixed in 4% PFA overnight at 4°C and were further thinned down manually using a scalpel until they were approximately 200 µm in thickness. The tissues were permeabilized with PBS containing 0.3 % Tween (PBST) overnight at 4°C and were blocked with 10% horse serum (in 0.3% PBST) for 1 hour at room temperature prior to overnight primary antibody staining at 4°C. The samples were washed for 15 minutes in 0.1 % PBST three times and were stained with the appropriate species-specific secondary antibodies for 2 hours at room temperature.

### High Speed Video Microscopy

Dissected mouse brains were sectioned in ice-cold PBS using a Leica VT1000S vibratome at 100 µm thickness. Coronal sections at bregma anterior/posterior (A/P) +0.30-0.40 mm were transferred in cell-imaging chambers containing pre-warmed serum-free Dulbecco’s Modified Eagle Medium (DMEM) and videos of the cilia beating were recorded on a live cell-configured inverted microscope attached to a high-speed video camera (Axio Observer Z1 platform equipped with ApoTome.2, Zeiss, Oberkochen, Germany) using the 40x objective at a stable temperature of 37°C and supplied with 5% CO_2_. The videos were recorded at a frame rate of 360 frames per second with a duration of at least 720 frames using the CoreView (v2.2.0.9) software (IO Industries, London, ON, Canada). The cilia beating frequency for each video was measured using the Fast Fourier transform (FFT) algorithm in Matlab (R2019a, Mathworks, Natick, MA). A cutoff of greater or equal to 2 Hz frequencies was included in the generated heatmap of each frame and the average cilia beating frequency (CBF) of each video was calculated as previously described (Oltean et al., 2018).

### Cell cultures

Wild type and *SHP2* knockout Human Embryonic Kidney 293 (HEK) cells were cultured in DMEM supplemented with 10% fetal calf serum (FCS) (Zhu et al., 2022). Transfection of the cells with pCDNA3.1. vectors expressing wild type SHP2, SHP2^E76K^, SHP2^C459S^ or SHP2^E76K/C459S^ was performed using Polyethyleneimine (PEI) STAR™ (7854, Tocris). The cells were starved in serum-free DMEM overnight prior to cytokine stimulation. Treatment of the cells with either Leukemia Inhibitory Factor (LIF) (200 ng/ml), Fibroblast Growth Factor 2 (FGF) (100 ng/ml) or Epidermal Growth Factor (EGF) (200 ng/ml) was performed 10 minutes prior to harvesting.

### Western blot

Dissected lateral walls or cultured cells were lysed in in CelLytic-M lysis buffer (C2978, Sigma) containing a proteinase inhibitor cocktail (PI78440, Thermo Fisher). The lysates were sonicated and then centrifuged at 12,000 g for 10 min to remove residual debris. The supernatants were collected and loading buffer containing anionic detergent Sodium Dodecyl Sulfate (SDS) and disulfide bond reducing agent, 2-mercaptoethaonl, was added to the samples. The mixture was boiled at 95°C for 5 min and equal amounts of total protein were loaded on an SDS-Page gel. The proteins were transferred for the gel to Polyvinylidene Difluoride (PVDF) membranes and were subsequently stained according to standard protocols. The membranes were scanned using an Odyssey SA scanner (LICOR Biosciences, Lincoln, NE).

### Differential scanning fluorimetry (DSF)

Full-length SHP2 proteins (wild type, E76K, C459S, and E76K/C459S) were expressed and purified as described previously (van Vlimmeren et al., 2024). The thermostability of each protein was measured using differential scanning fluorimetry (DSF) on MicroAmp Fast Optical 96-well Reaction plates (Applied Biosystems, 4346906) at working volumes of 20 μL. Purified proteins were diluted in DSF buffer (20 mM HEPES pH 7.5, 50 mM NaCl, 0.4% DMSO) to 10 μM in a mixture that also contained SYPRO Orange Protein Gel Stain (Thermo Fisher, S-6650), diluted 200-fold from a 5000x stock to 25x as the working concentration. Melting curves were measured on an Applied Biosystems Step-One Plus RT-PCR thermocycler between 25 °C and 95 °C with a gradient of +0.5 °C per minute (excitation: 472 nm; emission: 570 nm). Fluorescence reads under each temperature were analyzed using DSFworld and melting temperatures were calculated with dRFU (Wu et al., 2024).

### Statistical analysis

The quantification of the Glial Fibrillary Acidic Protein (GFAP) relative expression area and the number of Neuronal Nuclei (NeuN) positive cells of the entire hippocampus was performed using ImageJ. Vertical cortical thickness from the base of the ventricular zone to the pia of the primary motor and primary somatosensory cortex was measured using ImageJ calibrated to 1 mm. The averages of 5 technical replicates were used for statistical analysis and at least three biological replicates were included for each group. To distinguish differences between wild type and mutant mice, the Mann-Whitney U Test and Kruskal-Wallis one-way ANOVA were used. Signal intensity from western blot images of the phosphorylated protein and total protein was quantified using ImageJ. The Ratios between the phosphorylated and unphosphorylated protein levels were used to test differences between wildtype and conditional knockout mice using Mann-Whitney U Test.

## Acknowledgments

The authors thank Gen-sheng Feng for *Shp2^flox^* mice and Zhong-Yin Zhang, and W. Andy Tao for HEK293 *SHP2* KO cell line and *SHP2*-expressing vectors. The work was supported by grants from NIH (EY017061, EY018868 and EY025933 to X.Z. and R35GM138014 to N.H.S). N.M. is supported by a Knights Templar Eye Foundation Career Starter Grant Award. The Columbia Ophthalmology Core Facility is supported by NIH Core grant 5P30EY019007 and unrestricted funds from Research to Prevent Blindness (RPB).

